# GREIN: An Interactive Web Platform for Reanalyzing GEO RNA-seq Data

**DOI:** 10.1101/326223

**Authors:** Naim Al Mahi, Mehdi Fazel Najafabadi, Marcin Pilarczyk, Michal Kouril, Mario Medvedovic

## Abstract

The vast amount of RNA-seq data deposited in Gene Expression Omnibus (GEO) and Sequence Read Archive (SRA) is still a grossly underutilized resource for biomedical research. To remove technical roadblocks for reusing these data, we have developed a web-application GREIN (GEO RNA-seq Experiments Interactive Navigator) which provides user-friendly interfaces to manipulate and analyze GEO RNA-seq data. GREIN is powered by the back-end computational pipeline for uniform processing of RNA-seq data and the large number (>6,500) of already processed datasets. The front-end user interfaces provide a wealth of user-analytics options including sub-setting and downloading processed data, interactive visualization, statistical power analyses, construction of differential gene expression signatures and their comprehensive functional characterization, and connectivity analysis with LINCS L1000 data. The combination of the massive amount of back-end data and front-end analytics options driven by user-friendly interfaces makes GREIN a unique open-source resource for re-using GEO RNA-seq data. GREIN is accessible at: https://shiny.ilincs.org/grein, the source code at: https://github.com/uc-bd2k/grein, and the Docker container at: https://hub.docker.com/r/ucbd2k/grein.

## Introduction

Depositing RNA-seq datasets in Gene Expression Omnibus (GEO)^1^ and Sequence Read Archive (SRA)^2^ repositories ensures the reproducibility of published studies and facilitates its reuse. Re-analysis of these data can lead to novel scientific insights^3^ and it has been routinely used to inform the design of new studies^4^. However, reuse of GEO RNA-seq data is made difficult by the complexity of the processing protocols^5^ and analytical tools^6^ which are often inaccessible to biomedical scientists not specializing in bioinformatics.

Recent efforts at re-processing GEO/SRA RNA-seq data alleviate this problem by providing access to large number of processed and per-transcript summarized RNA-seq datasets which significantly simplifies its use^7-9^. Other resources provide access and analysis tools for specific datasets^10,11^. While these platforms are extremely useful, they do not support additional functionalities for downstream analyses. For example, exploratory data analysis, differential expression analysis with batch effect adjustment, or statistical power analysis (Table 1). Therefore, open-source user-friendly tools with comprehensive analytical toolbox for re-analysis of public RNA-seq data are still lacking. We address this problem by developing and deploying GEO RNA-seq Experiments Interactive Navigator (GREIN) web tool for analysis of GEO RNA-seq data. In addition to the rich repertoire of analysis tools, GREIN provides access to more than 6,500 uniformly processed human, mouse, and rat GEO RNA-seq datasets with >215,000 samples that are ready for analysis. These datasets were retrieved from GEO and uniformly reprocessed by the back-end GEO RNA-seq experiments processing pipeline (GREP2). The pipeline also curates metadata for each of the datasets and annotates each sample with biomedical ontologies provided by MetaSRA^12^. As the number of new studies are included in GEO, more datasets are processed and added to GREIN on a regular basis. Apart from the preprocessed datasets, GREIN also facilitates user requested processing of GEO RNA-seq datasets on-the-fly (Table 1). We also release GREP2 as an R^13^ package and GREIN as a Docker^14^ container for easy local deployment of the complete infrastructure which can be used to reproduce GREIN results off-line.

**Table 1.** Comparison of different RNA-seq data resources. All these resources provide access to processed RNA-seq data, however most of them do not provide interactive interfaces for further manipulation and analyses of the datasets.

| Comparison | RNA-seq processed data resources |  |  |  |  |
| --- | --- | --- | --- | --- | --- |
|  | GREIN | Recount2<sup>7</sup> | ARCHS4<sup>8</sup> | Toil<sup>9</sup> | Expression Atlas<sup>10</sup> |
| Data availability | Gene and transcript | Gene and transcript | Gene and transcript | Transcript | Gene |
| Quantification type | Read mapping (Salmon) | Read alignment (Rail-RNA) | Read mapping (Kallisto) | Read mapping (Kallisto) | Read alignment (Tophat) |
| Interactive exploratory analysis | Yes | No | No | No | No |
| QC report | Yes | No | No | Yes | Yes |
| Power analysis | Yes | No | No | No | No |
| Differential expression analysis on-the-fly | Yes | No | No | No | No |
| Enrichment analysis for differential expression signature | Yes | No | No | No | Yes |
| Pipeline availability | R package and Docker container | R package and Docker container | Docker container | Docker container | NA |
| Process data on-the-fly | Yes | No | No | No | No |
| Ontological annotations of samples | Yes | No | No | No | No |

## Results

The conceptual outline of GREIN is showed in Fig. 1. Individual RNA-seq datasets are processed by the GREP2 pipeline and stored locally as R Expression Sets. User can access and analyze preprocessed datasets via GREIN graphical user interface (GUI) or submit for processing datasets that have not yet been processed. GUI-driven workflows facilitate examination and visualization of data, statistical analysis, transcriptional signature construction, and systems biology interpretation of differentially expressed (DE) genes. Both GREIN and the back-end pipeline (GREP2) are written in R and released as Docker container and R package respectively. Graphical user interfaces for GREIN are implemented in Shiny^15^, a web framework for building dynamic web applications in R. The web instances at https://shiny.ilincs.org/grein is deployed via robust docker swarm of load-balanced of Shiny servers. The complete GREIN infrastructure, including processing pipeline is deployed via Docker containers.

**Figure 1.**
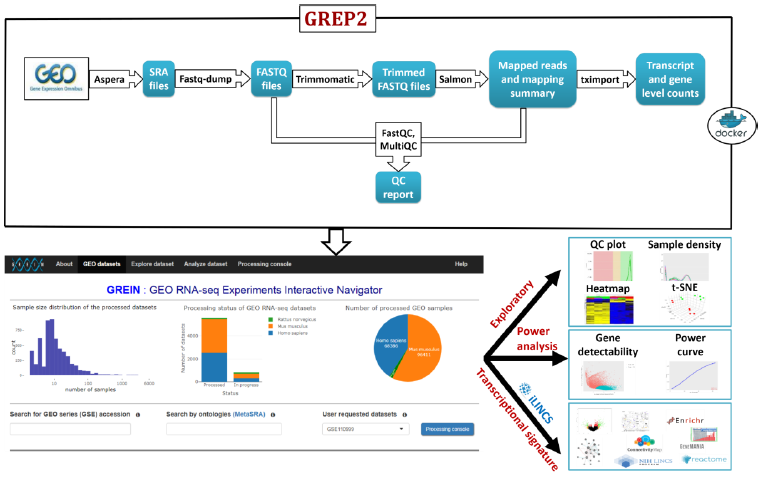
Schematic workflow of GREP2, web interface and outputs of GREIN. GEO datasets are systematically processed using GREP2 pipeline and stored within the back-end dataset library. GUI-driven GREIN workflows facilitate comprehensive analysis and visualization of processed datasets.

User friendly GUI driven workflows in GREIN facilitate typical reuse scenarios for RNA-seq data such as examination of quality control measures and visualization of expression patterns in the whole dataset, sample size and power analysis for the purpose of informing experimental design of future studies, statistical differential gene expression, gene list enrichment, and network analysis. Besides standard two-group comparison, the differential gene expression analysis module also supports fitting of a generalized linear model that accounts for covariates or batch-effects. The interactive visualization and exploration tools implemented include cluster analysis, interactive heatmaps, principal component analysis (PCA), t-distributed stochastic neighbor embedding (t-SNE), etc. (Supplementary Table S1). User can also search for ontological annotations of human RNA-seq samples and datasets provided by the MetaSRA project^12^. Each processed human RNA-seq sample is labelled with MetaSRA mapping of biomedical ontologies including Disease Ontology, Cell Ontology, Experimental Factor Ontology, Cellosaurus, and Uberon. Biological interpretation of differential gene expressions is aided by direct links to other online tools for performing typical post-hoc analyses such as the gene list and pathway enrichment analysis and the network analysis of differentially expressed (DE) genes. The connection to these analytical web services is implemented by submitting the differential gene expression signature (i.e., the list of average changes in gene expression and associated p-values for all/up/down regulated genes analyzed) to iLINCS^16^ (Integrative LINCS). iLINCS also provides the signatures connectivity analysis for recently released Connectivity Map L1000 signatures^17^. Detailed step-by-step instructions about GREIN analysis workflows are provided in the Supplementary Material and ‘Help’ section in GREIN.

## Key functionalities

### Search or submit for processing

User can either search for an already processed GEO data set in the ‘*Search for GEO series (GSE) accession*’ box or submit a dataset for processing if the dataset is not already processed (Supplementary Fig. S2). At this point in time, the vast majority of GEO human, mouse and rat RNA-seq datasets has been preprocessed and the user-submission of GEO datasets for processing will be required only occasionally. User can check the processing status of the requested dataset in the ‘*Processing console*’ tab (Supplementary Fig. S3). Other search options include keyword search through metadata of the datasets and search samples through biomedical ontologies via MetaSRA ontological annotations.

### Explore dataset

GREIN allows access to both raw and normalized (counts per million and transcript per million) gene and transcript level data. GREIN comes with several interactive and customizable tools to visualize expression patterns such as interactive heatmaps of clustered genes and samples, density plots for all or a subset of samples, between and within group variability analysis through 2D and 3D dimensionality reduction analyses and visualizations such as PCA and t-SNE (Fig. 2). User can also visualize expression profile of each gene separately (Supplementary Fig. S6). Comprehensive quality control (QC) report of raw sequence data and sequence mapping (Supplementary Fig. S7) provides read mapping statistics and quality scores of the sequence data files for each sample.

**Figure 2.**
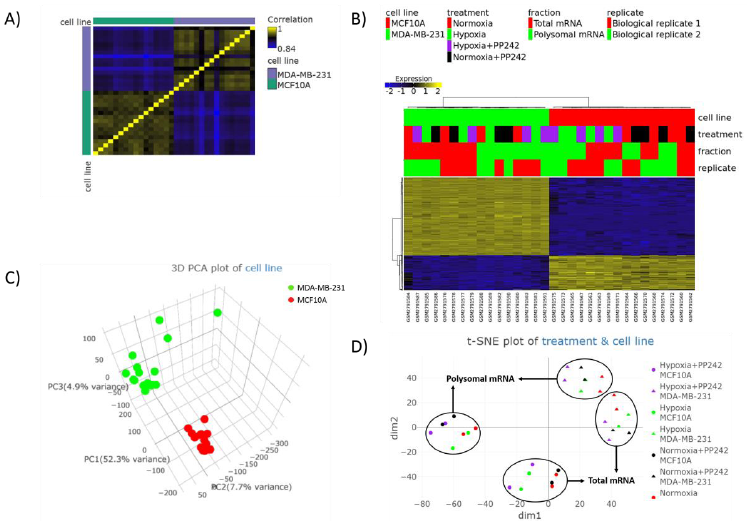
Exploratory analysis plots in GREIN. (**A**) Correlation heatmap shows a higher correlation within cell lines and low correlation between cell lines. Generally high correlations within each cell line indicate high quality of transcriptional profiles. (**B**) Hierarchical clustering based on Pearson correlation of top 500 most variable genes based on median absolute deviation as the variability measure. Data is normalized and centered to the mean. (**C**) Three-dimensional principal component analysis plot of the cell lines. (**D**) Two-dimensional t-SNE plot of treatment condition and cell line shows clear separation of the cell lines, and then the RNA fractions indicating.

### Statistical power analysis

The power analysis module in GREIN facilitates calculation and visualization of statistical power of detecting differentially expressed genes in future studies utilizing similar biological samples. Estimating appropriate sample size for future studies with similar biological samples is often the key motivating factor in re-analysis of RNA-seq data. Power analysis also facilitates the post-hoc analysis of false negative rates in the current dataset. The lack of statistical power and differences in statistical power between genes can produce false negative results leading to wrong conclusions^18^. The ‘*Power curve*’ segment provide power estimates for different number of samples based on a single gene (Fig. 3A). User can modify the default values of the parameters. The ‘*Detectability of genes*’ plot visualizes power estimate of each of the genes based on the selected groups and gene-wise dispersion (Fig. 3B). Mean coverage of the genes are plotted against their biological variability and are displayed in two sets based on their detectability status (power≥0.8 and power<0.8).

**Figure 3.**
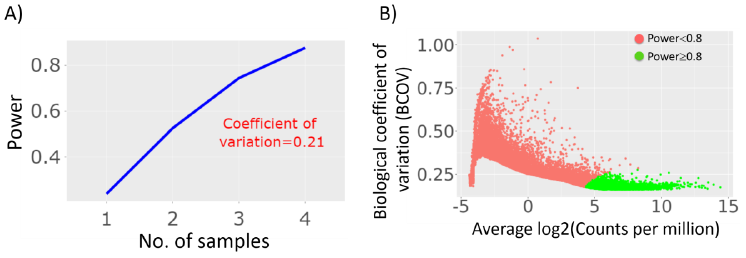
Power analysis for assessing transcriptional changes in non-malignant MCF10A cell line. (**A**) Single-gene based power estimates for different number of samples in each group with a minimal fold change of 2, statistical significance a=0.01, and a common dispersion estimate. (**B**) Gene-wise detectability on the log2CPM-BCOV plane with FDR≤0.1 and two samples in each group.

### Differential gene expression

Creating and interpreting differential gene expression signature is a typical analysis scenario in RNA-seq experiments. With GREIN, user can create a signature by comparing gene expression between two groups of samples with or without adjustments for experimental covariates or batch effects. GREIN can handle complex experimental designs by providing the flexibility of rearranging groups and sub-groups or selecting specific samples. Differential expression signature can be visualized via interactive graphics that include heatmap of top differentially deregulated genes (Supplementary Figure S21) ranked by false discovery rate (FDR), log fold change vs. log average expression (MA) plot (Supplementary Figure S22), and gene detectability plot (Supplementary Figure S23). Differential expression signature, with or without accounting for potentially false negative results, can be directly exported to iLINCS for enrichment and connectivity analysis.

## Use case: Analysis of transcriptional and translational regulation of hypoxia in non-malignant breast epithelial and triple-negative breast cancer cell lines

We demonstrate the usage of GREIN by re-analyzing a recently published GEO RNA-seq data (GSE104193). Sesé *et al*.^19^ examined the transcriptional and translational regulation of hormone-refractory triple-negative breast cancer (TNBC) subtype under a combination of hypoxia and mTOR (mechanistic target of rapamycin) inhibitor treatment. In particular, the authors analyzed the expression profiles of TNBC (MDA-MB-231) and non-malignant breast epithelial (MCF10A) cells exposed to normoxic (21% O_2_) and hypoxic (0.5% O_2_) conditions and/or treated with an mTORC1 and −2 inhibitor PP242. Each of the samples were sequenced for total (T) and polysome-bound (P) mRNA. The dataset contains 32 samples, representing two biological replicates for each combination of cell line, oxygen level, treatment status, and mRNA fraction.

Exploratory analysis of the processed dataset in GREIN (Fig. 2) shows that the strongest source of variation in between samples comes from differences between the two cell lines. This is re-enforced by the correlation analysis of full expression profiles (Fig. 2A), the hierarchical clustering of top 500 highly variable genes based on median absolute deviation (Fig. 2B), 3D PCA plot of the samples (Fig. 2C), and the 2D t-SNE plot (Fig. 2D). Furthermore, high correlations between expression profiles for the same cell line (Fig. 2A) indicates good signal-to-noise in the gene expression measurements. The additional substructure of data indicated by the 2D t-SNE plot has been examined by painting samples according to different attributes (Supplementary Fig. S1). This analysis revealed that separations within each cell line are induced by different mRNA fractions and then differences between experimental conditions.

Next, we used GREIN to perform statistical power analysis based on the pattern of biological variability observed in this dataset. We considered transcriptional profiles of each cell line exposed to hypoxia and treated with or without PP242 which leads to four comparisons. Assuming an expression difference of at least two-fold between the groups, at the statistical significance of α=.01, and with only two replicates in each group, statistical power of a gene to be detected as differentially expressed is below 0.55 in all the comparisons (Table 2). Our analysis indicates that one would need 4 replicates per group to achieve 80% power detecting two-fold change in expression (Table 2 and Fig. 3A). In a typical RNA-seq experiment, a sequencing depth of 20-30 million is sufficient to quantify gene expression for almost all genes^4,20^ which is also evident in this dataset. We also evaluated statistical power of each gene to be detected as differentially expressed from the ‘*Detectability of genes*’ plot. Average log of counts per million (CPM) values of the genes were plotted against gene-wise biological coefficient of variation (BCOV) and power was calculated for the corresponding genes (Fig. 3B). A controlled false discovery rate of 0.05 and expected percentage of true positives of 10% was used to estimate statistical significance. We define a gene to be detectable as differentially expressed in hypoxic condition if its power is 0.8 or above. As expected, there exists an inverse relationship between BCOV and power (Fig. 3B). Also, power to detect differential expression of a gene increases with a higher log CPM or effect size.

**Table 2.**
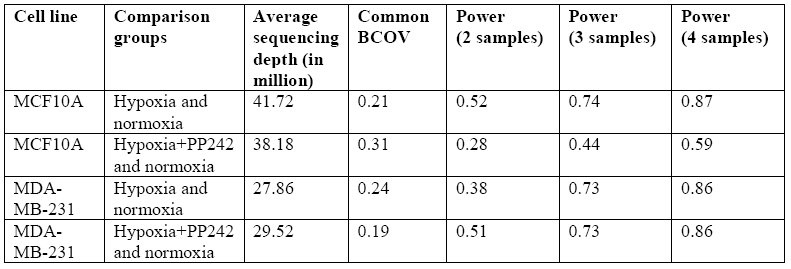
Statistical power analysis to assess transcriptional changes in malignant and non-malignant cell lines. With a minimal fold change of 2 between the groups and statistical significance at a=.01, all the comparisons are under-powered with two samples in each group. However, power increases as we increase the sample size.

| Cell line | Comparison groups | Average sequencing depth (in million) | Common BCOV | Power (2 samples) | Power (3 samples) | Power (4 samples) |
| --- | --- | --- | --- | --- | --- | --- |
| MCF10A | Hypoxia and normoxia | 41.72 | 0.21 | 0.52 | 0.74 | 0.87 |
| MCF10A | Hypoxia+PP242 and normoxia | 38.18 | 0.31 | 0.28 | 0.44 | 0.59 |
| MDA-MB-231 | Hypoxia and normoxia | 27.86 | 0.24 | 0.38 | 0.73 | 0.86 |
| MDA-MB-231 | Hypoxia+PP242 and normoxia | 29.52 | 0.19 | 0.51 | 0.73 | 0.86 |

One of the goals of the study was to analyze transcriptional changes in hypoxic and normoxic conditions with and without PP242 treatment in both MCF10A and MDA-MB-231 cell lines. We created transcriptional signatures of hypoxia and hypoxia+PP242 in total mRNA by differential expression analysis between hypoxia and hypoxia+PP242 samples respectively against the control samples while adjusting for batch effect by treating ‘*replicate*’ as a covariate, for each cell line separately. We found a higher number of genes differentially expressed (DE) in MCF10A cell lines compared to MDA-MB-231 in both hypoxia and hypoxia+PP242 (Fig. 4A) indicating that perhaps the tumor cell line is better equipped to deal with hypoxia. This analysis also showed that most non-differentially expressed genes are also not detectable, indicating that they may represent false negative results. This is in accordance with the power analysis showing that 4 samples per group would be needed to consistently identify differentially expressed genes with average BCOV. To identify lower expressed genes an even higher sample size would be required.

**Figure 4.**
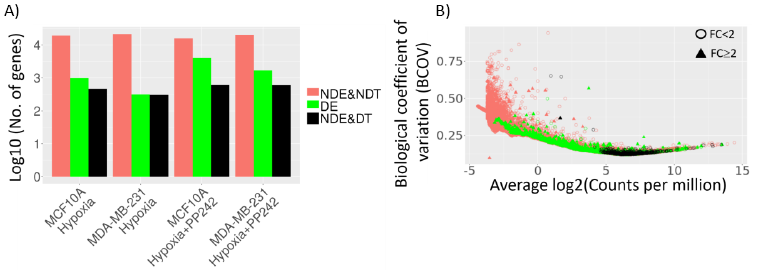
Differential expression and detectability of the genes. A) the number of genes (log10 scale) not differentially expressed and not detectable (NDE&NDT), differentially expressed (DE), and not differentially expressed but detectable (NDE&DT) in the comparisons with normoxia for total mRNA fraction. We call a gene detectable (DT) if its power≥0.8 and differentiable if FDR<0.05. B) The gene detectability plot for the first comparison (MCF10A and hypoxia) which visualizes the above-mentioned list of genes along with their respective fold changes (FC).

To interpret differentially expressed genes in terms of affected biological pathways, we submitted the differential gene expression signatures of hypoxia to online enrichment tools (DAVID^21^, ToppGene^22^, Enrichr^23^, and Reactome^24^) via iLINCS. The submitted signatures included a combined list of DE and NDE&DT genes representing likely true positive and true negatives. Genes were selected based on a cutoff of 0.7 and 0.01 for statistical power and FDR respectively. Fig. 5 illustrates the enrichment results obtained from ToppGene for the MCF10 hypoxia signature. Significantly enriched (FDR<0.05) top 10 gene ontology (GO) categories from ToppGene and DAVID functional annotation tool include response to hypoxia, response to decreased oxygen levels, angiogenesis, regulation of cell proliferation, oxidation-reduction process, and response to abiotic stimulus that are common in both cell lines (Supplementary Table S2 and Supplementary Table S3). Most of these categories are consistent with the original study. In addition, ToppGene suite identified hypoxia induced factor (HIF-1-alpha) transcription factor network that was activated in both cell lines (Supplementary Table S4 and Supplementary Table S5).

**Figure 5.**
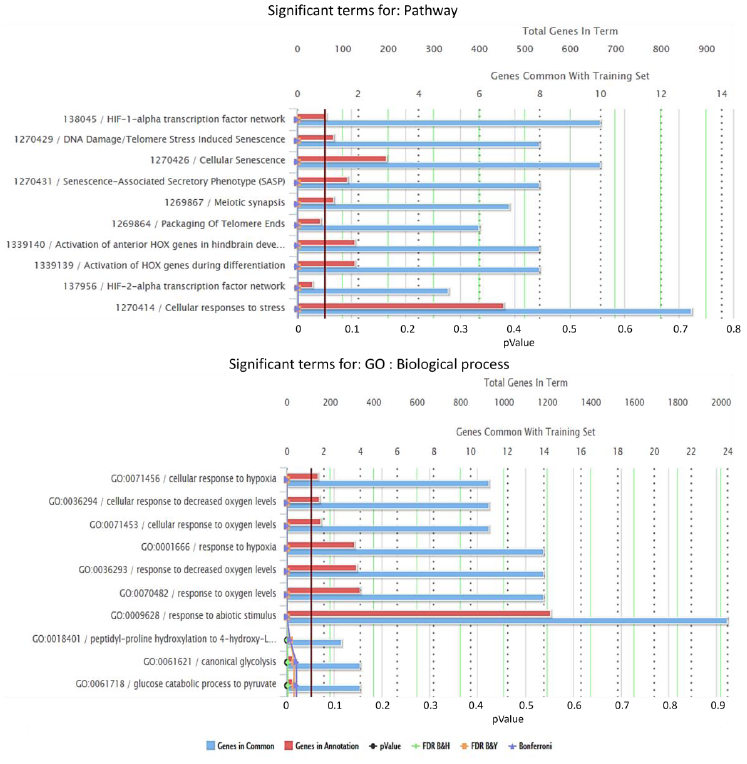
Snapshot of some of the significant pathway and gene ontology (GO) categories from ToppGene via iLINCS. These categories are found in the comparison between hypoxia and normoxia in MCF10A cell line using a combined list of DE and NDE&DT genes. The red vertical line is the selected cutoff of 0.05.

Finally, we utilized GREIN connection with iLINCS to “connect” the uploaded signature with LINCS^25^ consensus (CGS) gene knockdown signatures^17^. We found 3,727 LINCS consensus gene knockdown signatures that were significantly (pValue<0.05) connected with our uploaded signature. The target genes of top 100 connected signatures were selected for further enrichment analysis. We found cellular response to hypoxia and regulation of Hypoxia-inducible Factor (HIF) by oxygen in the list of top 10 activated pathways in both cell lines (Supplementary Table S6 and Supplementary Table S7). While this analysis yields similar enriched functional categories as the initial enrichment analysis, it complements the original analysis by implicating several target genes that are not differentially expressed although they are sufficiently highly expressed to be detectable according to our power analyses. Tying these two results together implicates these genes as potential higher level regulators of the response to hypoxia.

## Discussion

The combination of the access to a vast number of preprocessed mRNA expression data and the diversity of the analytical tools implemented makes GREIN a unique new resource for reuse of public domain RNA-seq data. GREIN not only removes technical barriers in reusing GEO RNA-seq data for biomedical scientists, but also facilitates easy assessment of the validity and reproducibility of the analysis results while uncovering new insights of the experiments. Our case-study analysis indicated that GREIN can be used for quickly reproducing results of the published studies as well as for gleaning additional insights that go beyond the original analysis. The complete analysis can be reproduced in less than 10 minutes. The web instance deployed on our server provides no-overhead use and requires no technical expertise. Open source R packages and the Docker container provide a flexible and transparent way for computationally savvy users to deploy the complete infrastructure locally with very little effort.

The GUI-driven analysis workflows implemented by GREIN covers a large portion of use cases for RNA-seq data analysis, making it the only tool that a scientist may need to meaningfully re-analyze GEO RNA-seq data. In addition, the power analysis workflow provides means to assess the statistical reasons for detecting or not-detecting specific differentially expressed genes. This kind of analyses is not common in the standard RNA-seq pipelines, but the results can be extremely useful when assessing false negative results. For example, when performing gene list enrichment analysis of differentially expressed genes, genes that do not meet a statistical significance cut-off are considered not differentially expressed. However, our power analysis indicates that the vast majority of these genes may simply be below detection limits for the available number of samples. GREIN allows user to distinguish between genes not detectable due to sample size limitations and/or their low expression levels, vs genes that are expressed at high enough expression levels but were not differentially expressed. This kind of resolution allows for more nuanced interpretation of analysis results. Off-line use of GREP2 and GREIN packages, as well as flexible export options enable additional analyses of GREIN-processed and pre-analyzed datasets.

## Methods

### GREIN back-end pipeline

To consistently process GEO RNA-seq datasets through a robust and uniform system, we have developed GEO RNA-seq experiments processing pipeline (GREP2) which is available as an R package in CRAN. Both GREP2 and GREIN are simultaneously running on different Docker containers. The whole processing workflow can be summarized in the following steps:

1. Obtain GEO series accession ID (series type: Expression profiling by high throughput sequencing) from GEO that contain at least two human, mouse, or rat RNA-seq samples. We then retrieve metadata for each GEO series accession using Bioconductor package GEOquery^26^. We also obtain metadata files from the Sequence Read Archive (SRA) to get corresponding run information and merge both the GEO and SRA metadata.
2. Download corresponding experiment run files from SRA using ‘ascp’ utility of Aspera Connect^27^ and convert them into FASTQ file format using NCBI SRA toolkit^28^. All the downloaded files are stored in the local repository until processed.
3. Run FastQC^29^ on each FASTQ file to generate QC reports and remove adapter sequences if necessary using Trimmomatic^30^.
4. Quantify transcript abundances by mapping reads to reference transcriptome using Salmon^31^.
5. Transcript level abundances are then then summarized to gene level using Bioconductor package tximport^32^. We use *lengthScaledTPM* option in the summarization step which gives estimated counts scaled up to library size while considering for transcript length. Gene annotation for Homo sapiens (GRCh38), Mus musculus (GRCm38), and Rattus norvegicus (Rnor_6.0) are obtained from Ensemble^33^ (release-91).
6. Compile FastQC reports and Salmon log files into a single interactive HTML report using MultiQC^34^.

### Power analysis

The power analysis in GREIN is performed using the Bioconductor package RNASeqPower^4^ which uses the following formula:

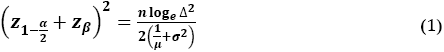

Where, α is the target false positive rate, β is the target false negative rate or 1-β is power, *n* is the sample size, Δ is the effect size, μ is the average sequencing depth, and σ is the biological coefficient of variation (BCOV) calculated as the square root of the dispersion. We use common dispersion and tagwise dispersion estimates from Bioconductor package edgeR^35^ for computing power of a single gene and multiple genes respectively.

Typically, thousands of genes are tested simultaneously for differential expression in RNA-seq experiments. Therefore, the above method for estimating power needs further adjustment to correct for multiple testing. Jung *et al*.^36^ derived an FDR correction formula and consequently the significance level (α^*^) to calculate sample size for microarray data in the following form:

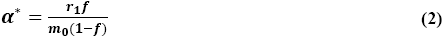

Where, *r*_1_ is the expected number of true positives in *m*_1_ rejected null hypotheses, *m*_0_ denote the number of genes for which null hypotheses are true, and *f* implies desired FDR level. Hence, to calculate power for each of the genes, we replace a with α^*^ in equation (1).

### Differential expression analysis

GREIN uses negative binomial generalized linear model as implemented in *edgeR* to find differentially expressed genes between sample groups. Data is normalized using trimmed mean of M-values (TMM) as implemented in edgeR. All the analyses are based on CPM values and genes are filtered at the onset with a cutoff of CPM>0 in *m* samples, where *m* is the minimum sample size in any of the groups. Besides two-group comparison, GREIN also supports adjustment for experimental covariates or batch effects. A design matrix is constructed with the selected variable and groups. We use gene-wise negative binomial generalized linear models with quasi-likelihood tests and gene-wise exact tests to calculate differential expression between groups with and without covariates respectively. P-values are adjusted for multiple testing correction using Benjamini-Hochberg method^37^. Interactive visualization of the differentially expressed genes is also available via heatmap of the top ranked genes, MA plot, and gene detectability plot.

## Acknowledgements

This work was supported by the National Institutes of Health (U54HL127624).

## Author Contributions

N.A.M. developed the pipeline and web application, M.M. conceived the project, supervised software development and data processing, M.M. and N.A.M. wrote the manuscript, M.F.N developed and maintain the Docker containers, M.P. and M.K. maintain the web server and implemented APIs for connecting with iLINCS. All authors reviewed the manuscript.

